# Towards understanding the mechanistic basis of a sex-limited color polymorphism

**DOI:** 10.64898/2026.05.02.722450

**Authors:** Tyra Westelius, Robin Pranter, Corin Stansfield, Natalia Zajac, Nathalie Feiner

## Abstract

The presence of multiple discrete color patterns within a species has captivated evolutionary biologists for more than a century, especially when such polymorphism is confined to one sex. The brown anole *Anolis sagrei* exhibits a female-limited polymorphism in dorsal patterning, which is controlled by allelic variation at the autosomal gene *CCDC170*. Here, we present and test a threshold model that can explain why the polymorphism is female-limited. We hypothesize that allelic variation at the *CCDC170* locus affects only female color pattern because this gene is co-expressed with its neighboring gene *ESR1*, highly expressed in female, but not male, embryos. By manipulating embryonic estradiol levels, we show that genetic males can be induced to express the polymorphism according to allelic variation at the *CCDC170* locus, which is naturally masked by low expression levels of this gene. Inversely, treating genetic females with fadrozole, which depletes estradiol, leads to monomorphic patterns irrespective of genotype, as for natural males. Using RT-qPCR, we demonstrate that these effects are accompanied by a direct influence of estradiol and fadrozole on gene expression levels of *CCDC170* and *ESR1*, thereby validating the threshold model. Our results suggest that the *CCDC170-ESR1-*locus is part of a mechanistic link between the morph-determining and the sex differentiation systems and provide a causal explanation for the developmental origin of a sex-limited color polymorphism.

## Introduction

The color patterns of fish, amphibians and reptiles arise through the spatial organization of pigment cell types (Kratochwil & Mallarino, 2023; Kuriyama et al., 2020). Lizards possess three different types of pigment cells: melanophores (containing black/brown pigments), xanthophores (containing yellow/orange/red pigments), and iridophores (containing guanin crystals). These cells originate in the neural crest and migrate throughout the early embryo before settling in the integument, with the resulting color pattern largely influenced by the absolute and relative number of different types of pigment cells, their organization and cellular content. Nevertheless, the occurrence of Mendelian inherited discrete color patterns within a species (i.e., pattern polymorphisms) suggests that transitions between morphs can be achieved by a single genetic change. Particularly interesting cases are polymorphisms that are limited to one sex since these imply a genetic and developmental link between the generation of color pattern and the mechanism of sex differentiation (Mank, 2023). For example, the morph-determining locus can be located on the sex chromosome of the heterogametic sex, which is the case for female-limited color polymorphism in the common cuckoo (Merondun et al., 2024) and color polymorphism in male guppies (Lindholm et al., 2004; Sandkam et al., 2021). In other cases, the morph-determining locus has been located to an autosomal genomic region containing genes involved in sex determination, most notably *doublesex* in insects (Hendrickx et al., 2022; Nishikawa et al., 2015). However, why allelic variation located close to a sex-determining gene can cause sex-limited color polymorphism is poorly understood. Uncovering these mechanisms can provide insights into how changes in gene regulation contribute to the origin and evolution of color patterns.

We have previously identified the genetic basis of a female-limited color pattern polymorphism in the brown anole lizard *Anolis sagrei* (Feiner et al., 2022), a species with genetic sex determination and well-differentiated X and Y chromosomes (Geneva et al., 2022). Females exhibit two discrete types of dorsal color patterns, referred to as diamond and chevron, while males invariably show a chevron pattern (Fig. 1A). We identified the gene *CCDC170*, encoding a protein containing coiled-coil domains, which has been linked to cell migration and polarization in humans (Jiang et al., 2017), as a strong candidate for causing the morph differences. Females with chevron patterns carry two copies of a recessive allele we termed the c-allele, while females with diamond patterns carry at least one copy of the dominant D-allele, which differs from the c-allele by 13 amino acid substitutions, likely producing a protein with different structure (Feiner et al., 2022). The gene *CCDC170* is located on the autosomal chromosome 1, 56 kb upstream of the *Estrogen Receptor 1* gene, which plays a crucial role in female sexual development across vertebrates (Bondesson et al., 2015). The *ESR1* gene encodes the Estrogen Receptor alpha (ERα), which binds estrogen hormones such as estradiol. Estrogen binding induces a conformational change that allows ERα to act as a transcription factor and interact with estrogen-response elements (EREs) near target genes. Thus, in the presence of estrogen hormones, cells that express *ESR1* can switch on or off a battery of genes regulated by EREs, including *ESR1* itself.

**Fig. 1.**
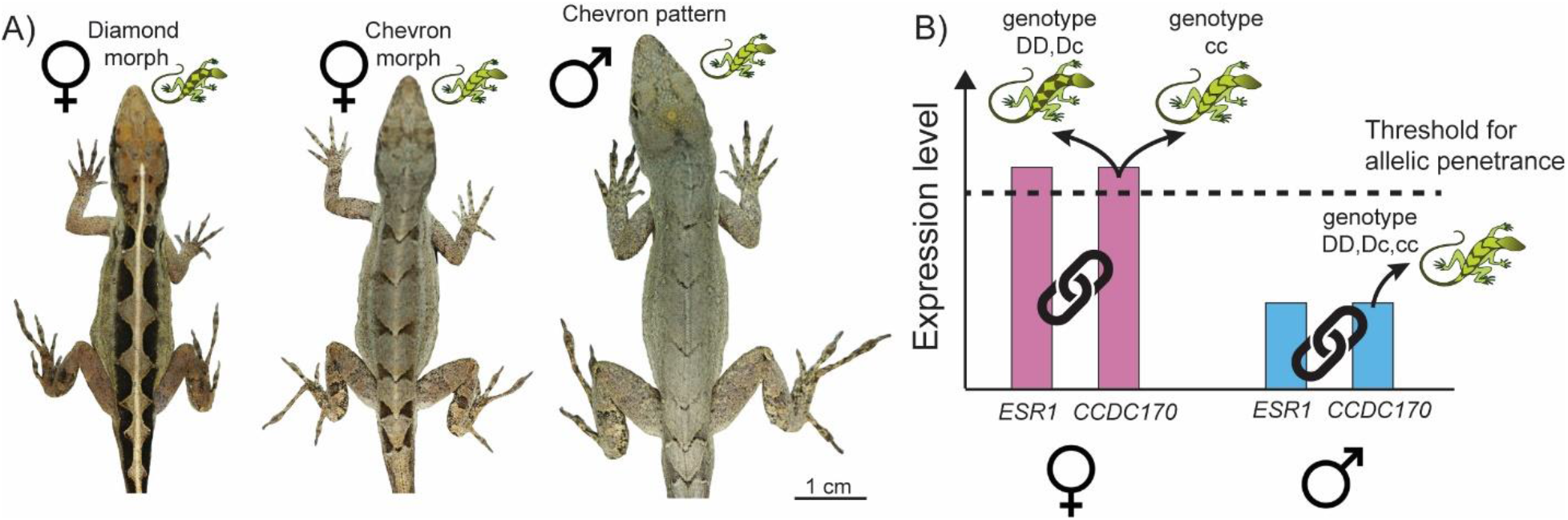
Dorsal color patterns of *A. sagrei* and threshold model for the effect of sex-biased gene expression on color morph determination. **A)** Representative photographs of the diamond and chevron morphs in females and the chevron pattern in males. **B)** Schematic drawing of hypothetical gene expression levels of *CCDC170* and *ESR1* in females (pink bars) and males (blue bars) and illustration of a threshold for allelic penetrance below which morph determination is not affected by allelic variation at the *CCDC170* locus. In contrast, females expressing *CCDC170* above a certain threshold develop diamond patterns if at least one D-allele is present at the *CCDC170* locus and chevron patterns if two c-alleles are present. Chain symbols illustrate co-expression between *CCDC170* and *ESR1*. Picture credit: Nathalie Feiner.

*ESR1* shows higher expression in female embryos of *A. sagrei*, both in secondary sexual organs (hemipenes that are present in embryos of both sexes) as well as in dorsal epithelia where color pattern formation is taking place (Feiner et al., 2022). We have previously shown that the *CCDC170* gene is co-expressed with the *ESR1* gene, potentially due to their close physical proximity, and that this co-expression is similar for both the c- and D-alleles. These observations led us to develop a threshold model that explains why the polymorphism is female-limited. We hypothesize that low expression of *CCDC170* in males prevents allelic penetrance, effectively ‘masking’ the pattern-determining locus and resulting in monomorphic males (Fig. 1B). In other words, an elevated expression of at least one dominant ‘D’ allele is required to generate a diamond pattern, and this is naturally restricted to females (Fig. 1B).

If this threshold model captures the relevant dynamics correctly, experimentally increasing *ESR1* expression levels should also upregulate *CCDC170*, causing genetic males that carry a D-allele to express the diamond morph. Similarly, experimentally decreasing *ESR1* expression levels should downregulate *CCDC170*, causing genetic females that carry a D-allele to express the chevron morph. Here, we experimentally manipulated embryos with estradiol and fadrozole, an inhibitor of the aromatase enzyme that converts testosterone to estradiol (Bhatnagar et al., 1990), to test these predictions. Administration of estradiol (i.e., the ERα ligand) is known to affect the expression of *ESR1* via a feedback loop. For example, studies in the Chinese soft-shelled turtle (Chen et al., 2022), a goldfish (Marlatt et al., 2008), two snake species (Katsu et al., 2010) and a wall lizard (Cardone et al., 1998) report increased expression levels of *ESR1* upon estradiol treatment. However, both up- and down-regulation of *ESR1* are reported in other species, sometimes with tissue-specific effects (Lung et al., 2020; Shupnik et al., 1989). For example, human cell lines exposed to exogenous estradiol tend to decrease expression of *ESR1* (e.g., Saceda et al., 1988). Thus, irrespective of how estradiol treatment of embryos changes *ESR1* expression, treatment with fadrozole should have the opposite effect. Indeed, it is well-established that estradiol and fadrozole administration during development are able to cause sex reversal in reptiles (Bull et al., 1988; Shine et al., 2007), including *A. sagrei* (Equinox, 2022).

Here we investigated if administration of estradiol and fadrozole to *A. sagrei* embryos affect both sex differentiation and color morph development according to the predictions outlined above. By measuring expression levels of *ESR1* and *CCDC170*, we obtained evidence consistent with the hypothesis that co-expression of these two genes represents a mechanistic explanation for female-limited color polymorphism.

## Materials & methods

### Animal husbandry

Breeding groups of *Anolis sagrei* were kept under conditions described in (Feiner et al., 2022; Feiner et al., 2020). Animals are the 2^nd^ and 3^rd^ generation of a breeding stock that was set up in 2016 from adult individuals wild-caught in Palm Coast, Florida, US. Eggs were collected every other day for two experiments as described below. In the first experiment, we administered estradiol or fadrozole to eggs to test its effect on the joint expression of sex and color pattern morph. In the second experiment, we used stressfully high and low incubation temperatures to test how incubation temperature affects variation within one morph.

### Treatment with estradiol or fadrozole

Eggs were divided into two groups and each group received treatment on day 0, 5 and 10. Treatment was administered by placing a 5 µl droplet onto the porous eggshell. In 2024, the control group received 95% ethanol, while the treatment group received 0.1 µg/µl 17β-estradiol (hereafter estradiol; for all reagents used in this study, see (Supplementary Table S1) in 95% ethanol freshly prepared on the day (Ehl et al., 2017). In 2025, the control group received 95% ethanol, while the treatment group received 0.6 µg/µl fadrozole in 95% ethanol (Ehl et al., 2017). Since the estradiol and fadrozole experiments were performed in different years, we analyzed them separately (see below). Eggs were incubated at 27°C in individual small plastic containers filled two-thirds with moist vermiculite (5:1 vermiculite:water volume ratio) and sealed with cling film. Embryos destined for RT-qPCR were dissected at day 14 in cold nuclease-free PBS and stored in RNA*later* (Qiagen) at −80°C. Embryos subjected to phenotyping (see below) were allowed to develop to a stage at which color patterns have formed (~28 days).

### Temperature incubation experiment

Eggs were divided into three groups and incubated at three temperatures: low (22°C), mid (27°C) or high (32°C). According to the literature, the low and high temperatures are below and above natural nest temperatures, respectively, and may therefore cause developmental stress (Pearson & Warner, 2016; Sanger et al., 2021; Sanger et al., 2018). Embryos were allowed to develop to a stage at which color patterns have formed (~70 days at low temperature, ~30 days at mid temperature and ~23 days at high temperature).

### RT-qPCR

Hemipenes and dorsal tissues were dissected in RNA*later* using fine needles and capsulotomy scissors and subjected to RNA extraction. Total RNA was extracted using the RNeasy Micro Kit (Qiagen) and reverse-transcribed into cDNA using SuperScript III (Invitrogen, Carlsbad, USA) following the instructions of the 3’-RACE System (Invitrogen, Carlsbad, USA; including RNase treatment of final cDNA). RT-qPCRs were conducted for three genes (*GAPDH* as reference gene for internal normalization and *CCDC170* and *ESR1* as target genes) using equimolar amounts of cDNA of each sample as template and each gene/sample combination was performed in three technical replicates. For further details on methodology, including primer sequences and PCR conditions, see Feiner et al. (2022).

### Phenotyping of color pattern and anatomical sex

Embryos that were allowed to develop to stages at which color patterns are clearly discernible were killed by transferring the eggs to −80°C. Dead embryos were dissected and pictures of their dorsal color patterns were taken. Chevron and diamond morphs show discrete differences and were scored by one observer (T.W.). Variation within the chevron morph was not quantified but we scored variation within the diamond morph that can be described as a continuum from a single longitudinal bar to a diamond pattern. We captured this variation using a sinuosity score (cf. Moon & Kamath, 2019) that is calculated as the geometric mean of the length of left and right diamond outline divided by the length of the midline of the dorsum between fore- and hind-limb insertions. A sinuosity score close to 1 indicates a bar-like pattern while an increasing score indicates a more diamond-like pattern. Anatomical sex was scored based on presence or absence of hemipenes.

### Genotyping and molecular sexing

Tail tissue was collected from each embryo and DNA was extracted using the DNeasy Blood & Tissue Kit (Qiagen, USA). We used two PCR-based genotyping assays to determine the *CCDC170* genotype and to determine the genetic sex following methods described in Feiner et al. (2022). In brief, the *CCDC170* genotype was determined by amplifying a 710-bp fragment of exon 2 and treating the PCR products with the restriction enzyme Taq I, whose recognition site is present in the D-, but not the c-allele. For molecular sexing, we used the primers AcarB previously developed by Gamble and Zarkower (2014), which target a male-specific fragment of 238 bp located on the Y chromosome.

### Statistical analyses

We conducted all statistical analyses in R (version 4.5.2; R Core Team, 2017). Differences in embryo survival, deviations from 1:1 sex ratios, and differences in the frequencies of morphs correctly predicted from genotypes were assessed using Fisher’s exact tests and chi-square tests, as appropriate. Differences in sinuosity scores were analyzed using Analysis of Variance, followed by *post hoc* pairwise comparisons using Tukey’s HSD. To assess the effects of treatment, sex, and their interaction on gene expression levels, we fitted linear models. When interaction terms were not significant *(P* > 0.3), they were removed and main effects were interpreted from the reduced model. When a significant interaction was detected, we conducted *post hoc* analyses using estimated marginal means implemented in the emmeans package (version 2.0.1), with pairwise comparisons to evaluate treatment effects within each sex.

### Genomic characterization of the CCDC170-ESR1 locus

To predict estrogen-response elements (EREs) in the vicinity of the *CCDC170-ESR1* genomic region and across the entire *Anolis sagrei* genome, we downloaded the consensus motif from Jaspar (https://jaspar.elixir.no/; accessed 29.01.2026; Matrix ID: MA0112.1). The DNA-binding domains of Erα receptors are identical between human, chicken, and alligator (Rice et al., 2017), suggesting a high level of conservation of ERα, and thus EREs, among tetrapods. We therefore performed motif discovery in the *A. sagrei* genome (ID) with this consensus motif using the command line version of FIMO v5.5.9 from MEME-SUITE. A statistical threshold of *P* < 0.01 was used for discovery of motif occurrence.

To assess if the *CCDC170* and *ESR1* genes are located in tandem also in other vertebrate taxa, we used OMA (https://omabrowser.org/) to examine the genomic neighborhood of the *CCDC170* gene across 139 vertebrate species belonging to the Hierarchical Orthologous Group HOG:E0789397.1a. Local gene-centric synteny in OMA indicated this collinearity is highly conserved across all vertebrates except in actinopterygian fish. We thus selected representative species per clade and conducted a collinearity analysis of the broader genomic region in 23 vertebrate species. Annotations were downloaded from NCBI RefSeq (Supplementary Table S2). Regions in the genome containing *CCDC170* together with neighboring genes were visualized in R using the R package gggenomes (v1.0.1).

## Results

### Treating developing embryos with estradiol and fadrozole affects sexual development

A total of 88 and 41 eggs were treated with estradiol and fadrozole, respectively, or a control treatment using 95% ethanol (control sample size: estradiol experiment, 69; fadrozole experiment, 35). Embryo survival was above 94% across all four treatment groups and did not differ between treatment and control groups (estradiol treated, 94.3%; estradiol control, 98.6%; Fisher’s exact test, *P* = 0.23; fadrozole treated, 95.1%; fadrozole control, 94.3%; *P* = 1). As previously reported in other taxa (Ehl et al., 2017; Equinox, 2022), the anatomical sex was affected by treatment with estradiol or fadrozole. At an embryonic stage close to hatching, males typically possess hemipenes that are clearly visible as paired structures (Fig. 2A).

**Fig. 2.**
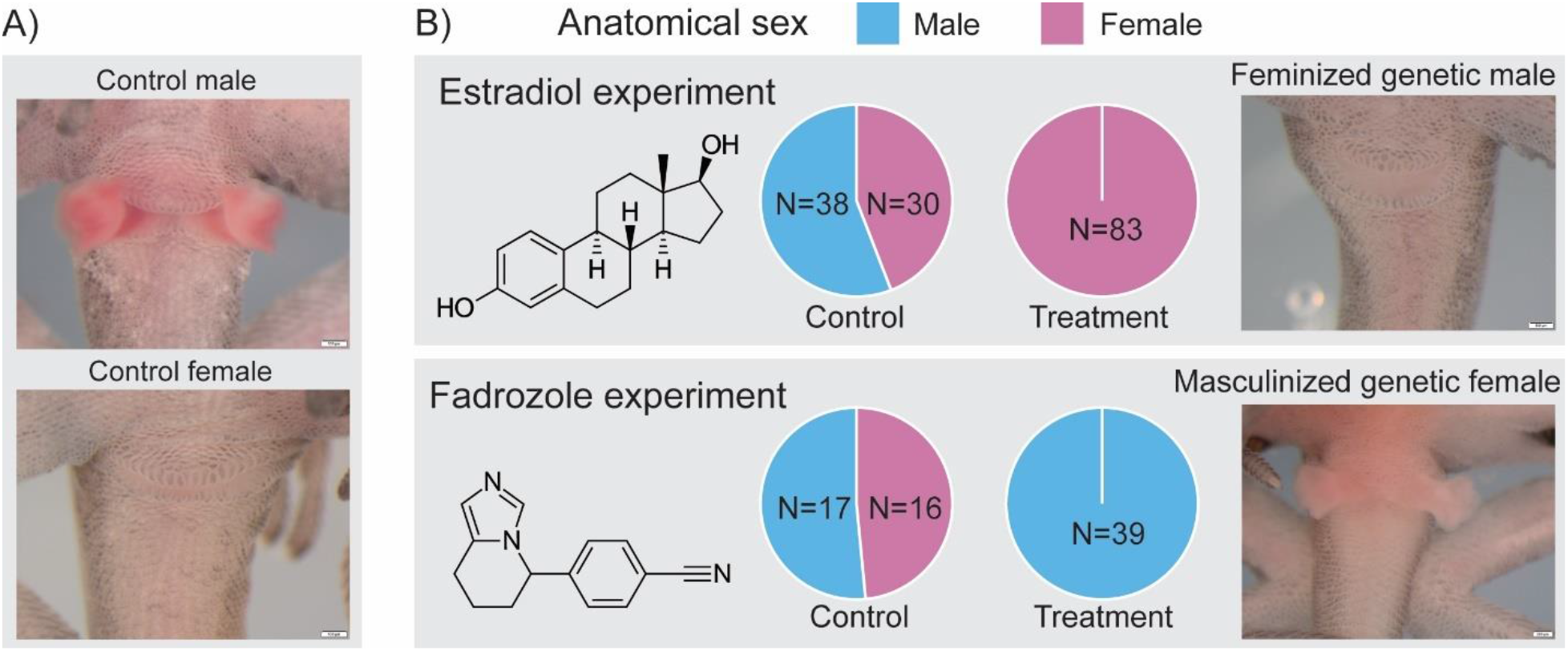
Effects of estradiol and fadrozole on anatomical sex. **A)** In control animals that were treated with ethanol, genetic males exhibit clearly visible hemipenes, while genetic females lack these structures at late embryonic stages. **B)** This characterization of the anatomical sex is affected by administration of estradiol (feminizing effect) or fadrozole (masculinizing effect). Pie charts present the number of anatomical males and females per treatment group (control or treatment) and experiment (estradiol or fadrozole). Images to the right show examples of the cloacal region of a feminized genetic male (estradiol treatment) and a masculinized genetic female (fadrozole treatment).

While hemipenes are also present in females at earlier stages of development, they have regressed and are not visible at the embryonic stage assessed here (Fig. 2A). Treating developing eggs with estradiol caused all embryos (N=83) to develop into anatomical females that lack visible hemipenes, while treatment with fadrozole caused all embryos (N=39) to develop into anatomical males with hemipenes (Fig. 2B). Molecular sexing verified that all four treatment groups showed the expected 1-to-1 genetic sex ratio (estradiol experiment, χ^2^_(1)_ = 0, *P* = 1; fadrozole experiment, χ^2^_(1)_ = 0.07, *P* = 0.80). Individuals were not raised to sexual maturity and it is therefore unknown to what extent primary sexual organs and sexual behavior are affected.

### Treatment with estradiol leads to upregulation of *ESR1* expression via a positive feedback loop

The consistent effect of estradiol and fadrozole on the anatomical sex before hatching may be due to their effect on the expression of the estradiol receptor, *ESR1*. Characterizing the genomic region of *ESR1*, we found that there are eight EREs in its vicinity, which is more than other genes in the *A. sagrei* genome (*P* = 0.040; Fig. 3A). Three of these EREs are directly upstream of the first exon. In addition, there were two EREs located downstream of *ESR1*, and two in the gene *CCDC170*, upstream of *ESR1*. This makes it likely that transcriptional regulation of *ESR1* is regulated by estradiol via a positive feedback loop, as has been shown for other vertebrates (see Introduction). For each treatment group, we performed RT-qPCR on hemipenes of embryos at embryonic stages 11-12 (staging based on Sanger et al., 2008) to quantify expression levels of *ESR1*. At this stage, both sexes possess hemipenes, but they are larger in males. Genetic sex had an overall significant effect on expression levels of *ESR1* in hemipenes, while the effect of treatment was negligible (Table 1, Fig. 3B). A *post hoc* test revealed that females in the control group showed a higher *ESR1* expression level in hemipenes than males in the control group (t = 2.339, df = 26, *P* = 0.027), but that this difference disappeared when embryos were treated with estradiol (t = 0.331, df = 26, *P* = 0.744). Treatment with fadrozole showed a similar effect in terms of removing sex-biased gene expression: while both treatment and genetic sex had a significant effect on the expression of *ESR1* in hemipenes (Table 1, Fig. 3B), a *post hoc* test showed that the difference between the sexes observed in the control groups disappears upon treatment with fadrozole (fadrozole-treated: t = 0.109, df = 18, *P* = 0.915; control: t = 4.409, df = 18, *P* = <0.001). Overall, these results indicate that treatment with estradiol upregulates the naturally low *ESR1* expression in male hemipenes to levels typical of females, while fadrozole downregulates the naturally high *ESR1* expression in female hemipenes to levels typical of males (Fig. 3B).

**Table 1.**
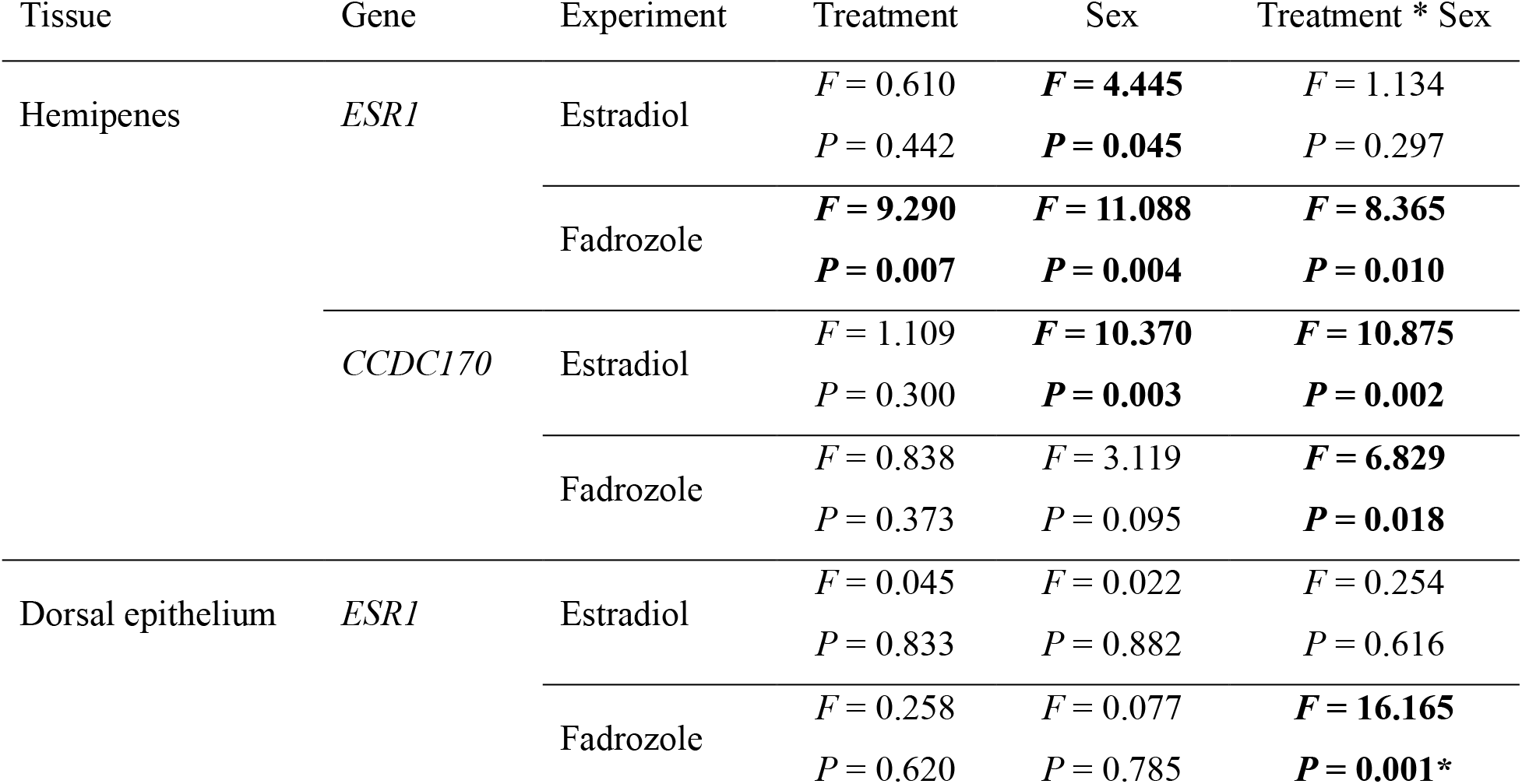
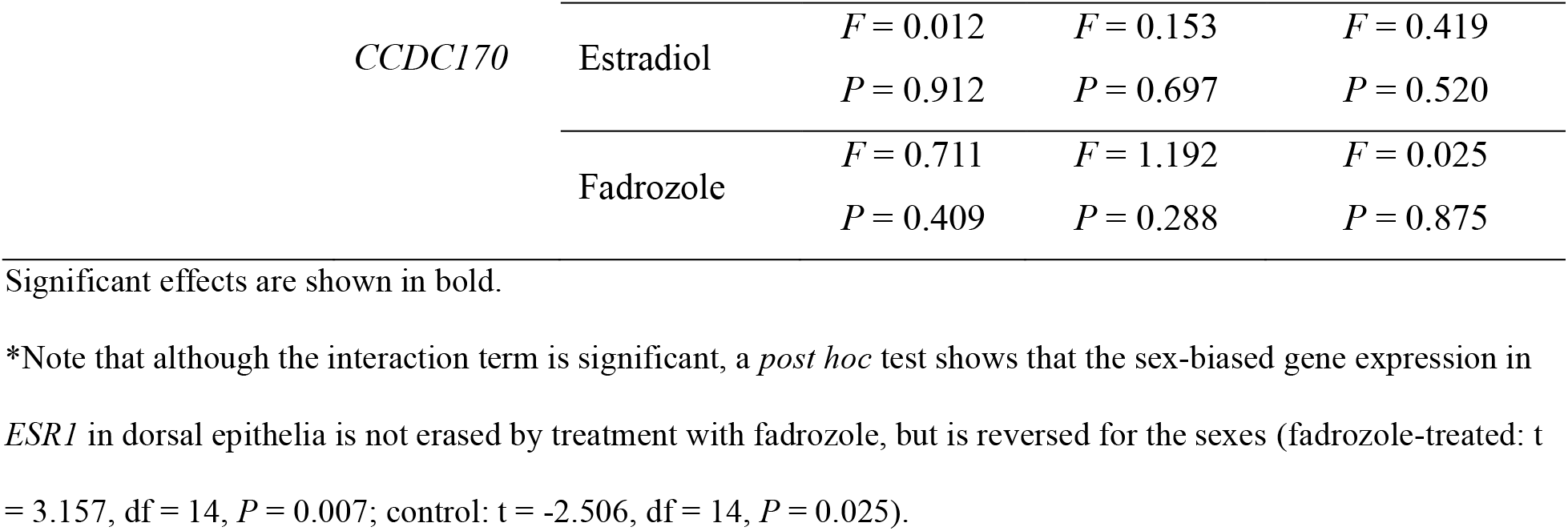
Effect of treatment of developing embryos with estradiol and fadrozole on expression levels of *ESR1* and *CCDC170* in hemipenes and dorsal epithelia.

**Fig. 3.**
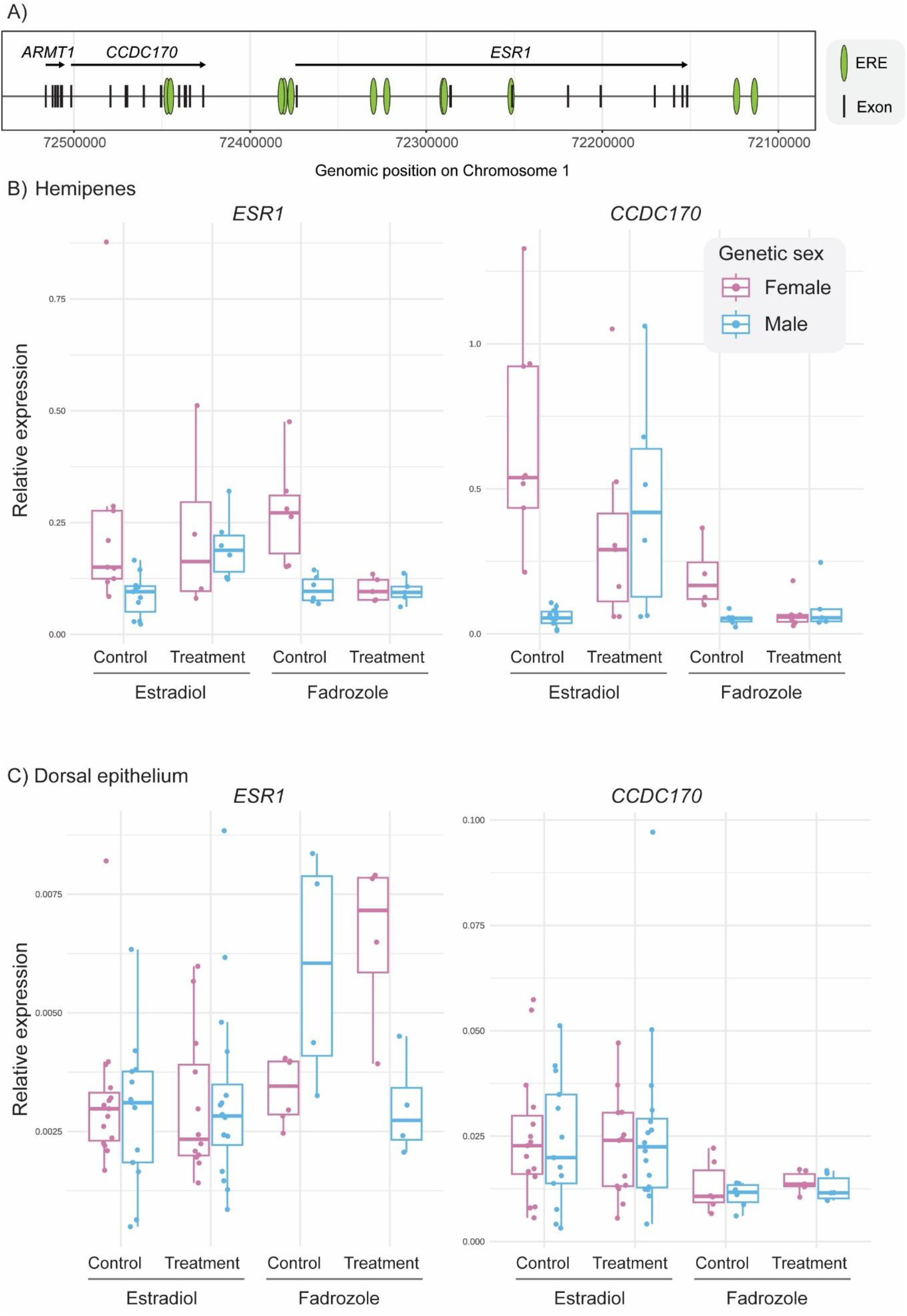
Effect of estradiol and fadrozole on the expression levels of *ESR1* and *CCDC170*. **A)** Predicted location of EREs in the genomic environment of *CCDC170* and *ESR1* are shown as green ellipses. **B)** and **C)** Relative expression of the two target genes (normalized using *GAPDH*) are plotted for hemipenes (panel B) and dorsal epithelia (panel C). Each datapoint presents the mean of three technical replicates. Boxes show interquartile ranges with middle line showing the median. Whiskers extend to 1.5 times the interquartile ranges.

Consistent with previous findings reporting co-expression between *ESR1* and the neighboring gene *CCDC170* (Feiner et al., 2022), we found that *CCDC170* expression levels respond in similar ways as *ESR1* in hemipenes upon estradiol and fadrozole treatment (Fig. 3C). While neither treatment had a significant main effect (Table 1, Fig. 3C), *post hoc* tests demonstrated that estradiol increases *CCDC170* expression in treated genetic males to levels typical of genetic females (estradiol-treated: t = −0.621, df = 31, *P* = 0.539; control: t = 4.567, df = 31, *P* = <0.001). *Vice versa*, fadrozole-treatment reduced the *CCDC170* expression in treated genetic females to levels typical of genetic males (fadrozole-treated: t = −0.500, df = 17, *P* = 0.624; control: t = 3.114, df = 17, *P* = 0.006; Fig. 3C). To test if the strong, sex-specific effect of estradiol and fadrozole on *ESR1* and *CCDC170* in developing hemipenes is also observed in dorsal epithelia, the tissue in which color patterns are formed, we performed the equivalent RT-qPCRs on dissected epithelia. We found that the overall, mean expression levels of *ESR1* and *CCDC170* were more than a magnitude lower in dorsal epithelia relative to hemipenes (mean expression levels: *ESR1*_Hemipenes_ = 0.165; *ESR1*_Dorsal Epithelium_ = 0.003; *CCDC170*_Hemipenes_ = 0.242; *CCDC170*_Dorsal Epithelium_ = 0.021). The only significant effect of sex and treatment was found for fadrozole, which appears to reverse sex-biased expression of *CCDC170* (Table 1).

Overall, we conclude that administration of estradiol leads to upregulation of *CCDC170* and *ESR1* via a positive feedback loop, while fadrozole has the opposite effect. However, these treatments appear to have limited quantitative effect on the naturally high expression of these genes in females, and the naturally low expression in males, resulting in an absence of sex-biased gene expression in estradiol- and fadrozole-treated embryos.

### Treatment with estradiol and fadrozole affect morph determination in a predictable way

Given that estradiol treatment increases, and fadrozole treatment decreases, *CCDC170* (and *ESR1*) expression levels, the threshold model predicts consistent effects on the expression of color morphs. Experimentally increasing *CCDC170* levels in genetic males using estradiol should ‘unmask’ the effects of allelic variation (c- or D-alleles) on color morph determination. In contrast, experimentally decreasing *CCDC170* levels in genetic females using fadrozole should ‘mask’ allelic variation at this morph determining locus, leading to the expression of the chevron morph, regardless of the allelic variation at the *CCDC170* locus.

In line with these predictions, we observed that all genetic males in the control treatment expressed the chevron color pattern, while treating genetic males with estradiol caused them to develop color patterns according to their *CCDC170* genotypes: all 34 estradiol-treated genetic males with DD or Dc genotypes developed a diamond morph, while 9 out of 10 estradiol-treated males with the cc genotype showed a chevron pattern (Fig. 4A). Thus, 43 out of 44 estradiol-treated genetic males developed a color morph that was predicted by their *CCDC170* genotypes, while this was only the case for 10 out of 36 animals in the control group where all were chevron morphs (Fisher’s exact test, *P* < 0.001). In contrast, treating genetic females with estradiol had no effect on the relationship between *CCDC170* genotypes and color morph, and genotype predicted the observed color morph in 31 out of 32 cases in the control group and 32 out of 39 cases in the estradiol-treatment group (Fisher’s exact test, *P* = 0.065). Thus, estradiol treatment ‘unmasked’ the effect of the *CCDC170* genotype on morph determination in genetic males, while fadrozole ‘masked’ the effect in females: all 16 genetic females that had a DD or Dc genotype treated with fadrozole developed a chevron phenotype (Fig. 4B). While *CCDC170* genotypes predicted the correct color morph in 14 out of 15 cases in the control group in genetic females, this was only the case for 2 out of 20 individuals in the fadrozole treatment group (Fisher’s exact test, *P* < 0.001). In contrast, fadrozole treatment had no effect on color morph in genetic males, with all embryos of both treatment and control groups expressing the chevron morph, regardless of their underlying *CCDC170* genotypes (Fisher’s exact test, *P* = 0.125). These findings are in agreement with the threshold model and suggest that the expression of *CCDC170* determines whether or not the allelic variation at this locus is manifested as color pattern morph determination.

**Fig. 4.**
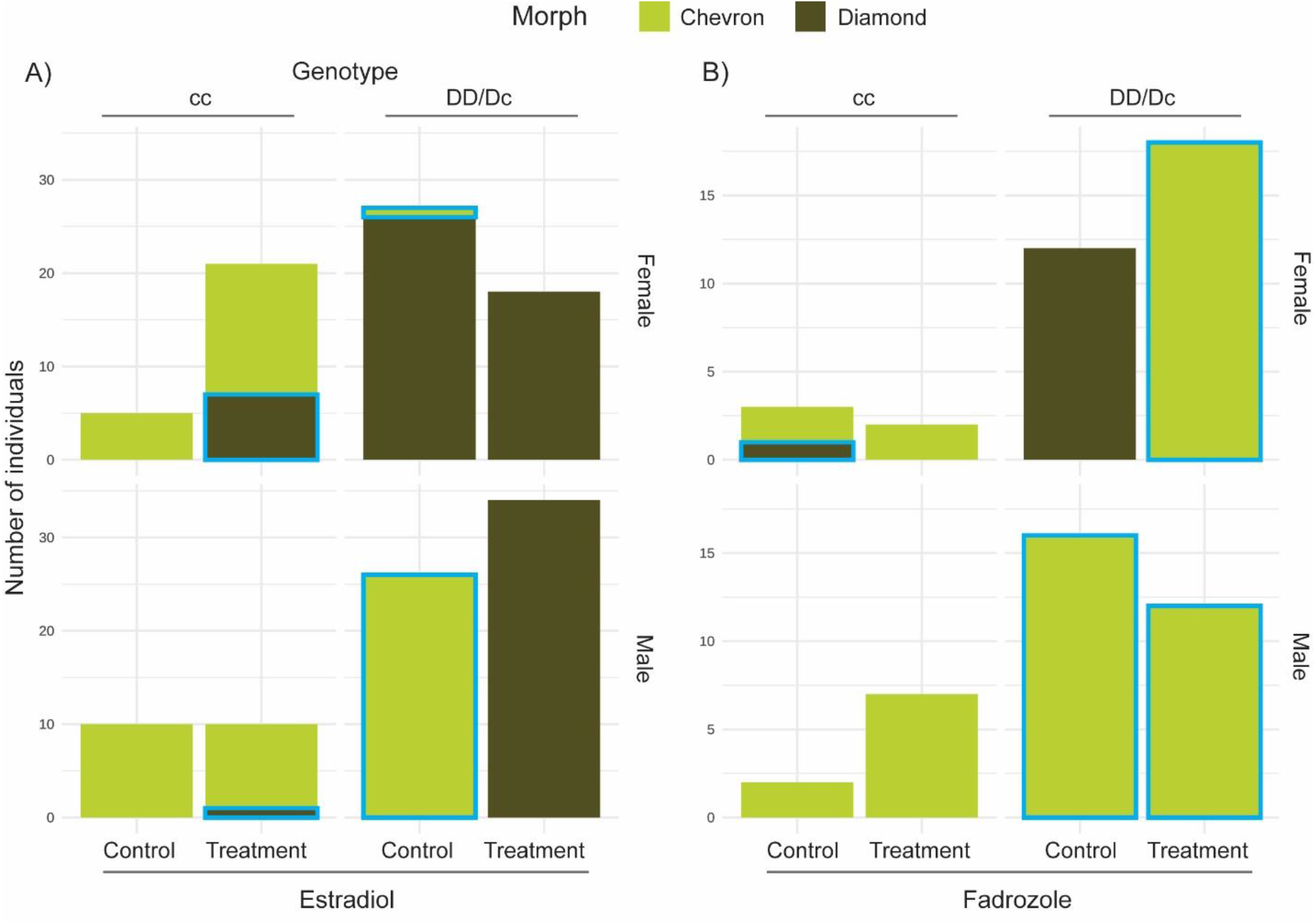
Effect of estradiol and fadrozole on color pattern morph in *A. sagrei*. **A)** Bar plots show the numbers of individuals showing chevron (green) and diamond (brown) morph for genetic females and males in the estradiol experiment treated either with estradiol (treatment) or ethanol (control). **B)** Same as in panel A, but for the fadrozole experiment. Blue outlines mark individuals where cc genotypes were not associated with chevron patterns or DD or Dc genotypes were not associated with diamond pattern (i.e., cases where genotype does not predict phenotype). Note that this is naturally the case for males with a DD or Dc genotype.

### Sex and environmental effects on quantitative aspects of the diamond pattern

After establishing that genetic males that possess at least one ‘D’ allele at the *CCDC170* locus develop a diamond pattern when treated with estradiol, we investigated whether these diamond patterns differ quantitatively from the naturally observed diamond patterns of females. We found that the sinuosity score was significantly different between males treated with estradiol and females treated with estradiol or control (*F* = 6.64; *P =* 0.002). A *post hoc* test showed that color patterns of males treated with estradiol are significantly more bar-like compared to females of the control treatment (*P* = 0.002). Estradiol-treated females also showed a trend towards a more bar-like pattern but this difference did not reach statistical significance (*P* = 0.128).

To test if this quantitative variation within the diamond morph reflects a disruption of optimal developmental conditions, we conducted an experiment in which we exposed developing embryos to stressfully low and high, in addition to optimal (here referred to as ‘mid’) incubation temperatures. In particular high incubation temperatures have been shown to lead to developmental stress (Sanger et al., 2021; Sanger et al., 2018). We found significant differences between the three temperature groups (*F* = 8.22; *P* < 0.001), with the high temperatures causing significantly more bar-like patterns with low sinuosity scores (low-high, *P* < 0.001; mid-high, *P* = 0.027). This suggests that high temperatures as well as hormone levels quantitatively influence dorsal pattern formation in *A. sagrei*, but that this effect does not ‘override’ the effect of D- and c-alleles on morph determination.

### Coupling between sex- and morph-determination at the genomic level

Our results suggest that morph determination is linked to sex differentiation through the genetic linkage between *CCDC170* and *ESR1*. Given that female-limited polymorphisms have been reported in other vertebrates (Dijkstra et al., 2007; Falk et al., 2022), we investigated if the collinearity between the two genes is broadly conserved. We find that across vertebrates, *CCDC170* and *ESR1* were located next to each other and in the same orientation in all ten investigated mammals, in four out of five reptiles (incl. birds), in both amphibians and in all three chondrichthyans (Fig. 6). Actinopterygian fish genomes did not preserve the linkage between *CCDC170* and *ESR1*. The overall high level of conservation in collinearity may indicate that the mechanisms for sex-linked color pattern morphs in *Anolis sagrei* may be broadly applicable to lizards and other vertebrates.

**Fig. 5.**
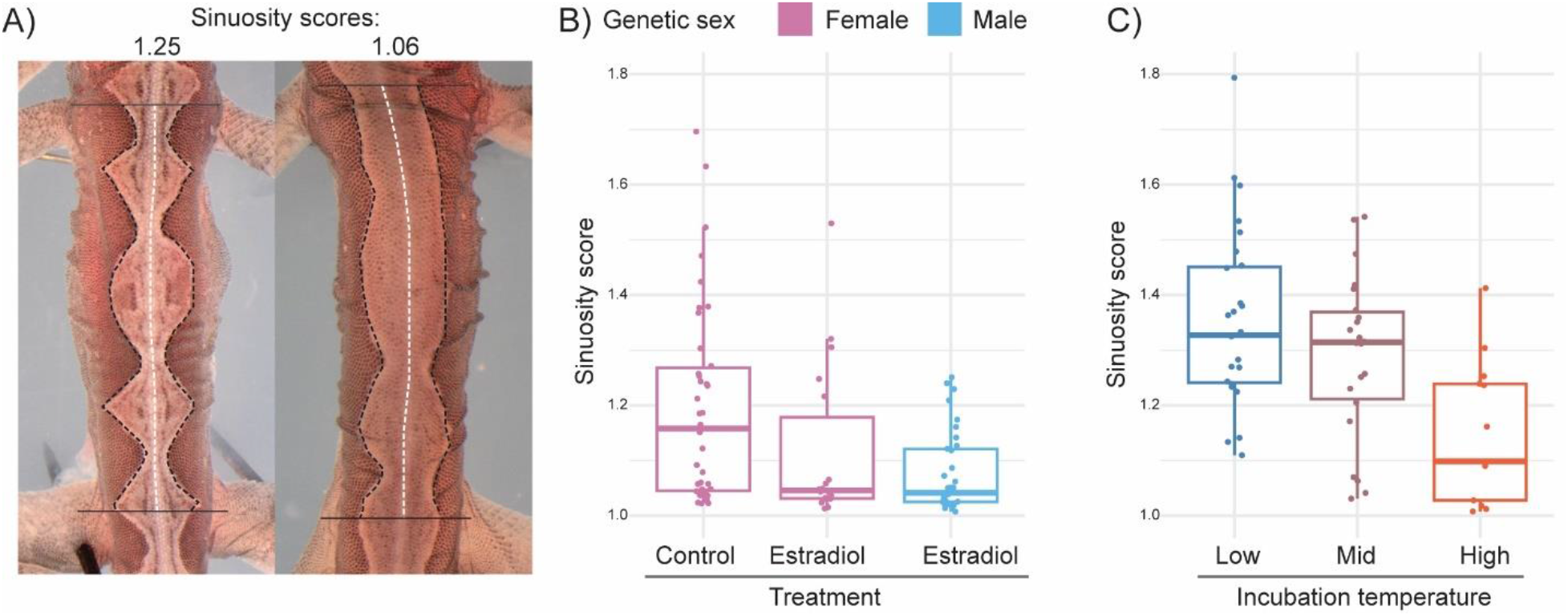
Effect of estradiol and incubation temperature on diamond pattern formation. **A)** Two diamond patterns exemplifying a diamond-like (left) and a more bar-like (right) phenotype. The sinuosity score is derived from the geometric mean of the length of left and right diamond outline (black dashed lines) divided by the length of the midline (white dashed line) between fore- and hind-limb insertions (marked by grey horizontal lines). **B)** Sinuosity scores of genetic females treated with ethanol (control; N = 38) or estradiol (N = 18) and genetic males treated with estradiol (N = 34). **C)** Sinuosity scores of untreated females that were incubated at low (N = 31), mid (N = 27) or high (N = 14) temperatures. Boxes show interquartile ranges with middle line showing the median. Whiskers extend to 1.5 times the interquartile ranges.

**Fig. 6.**
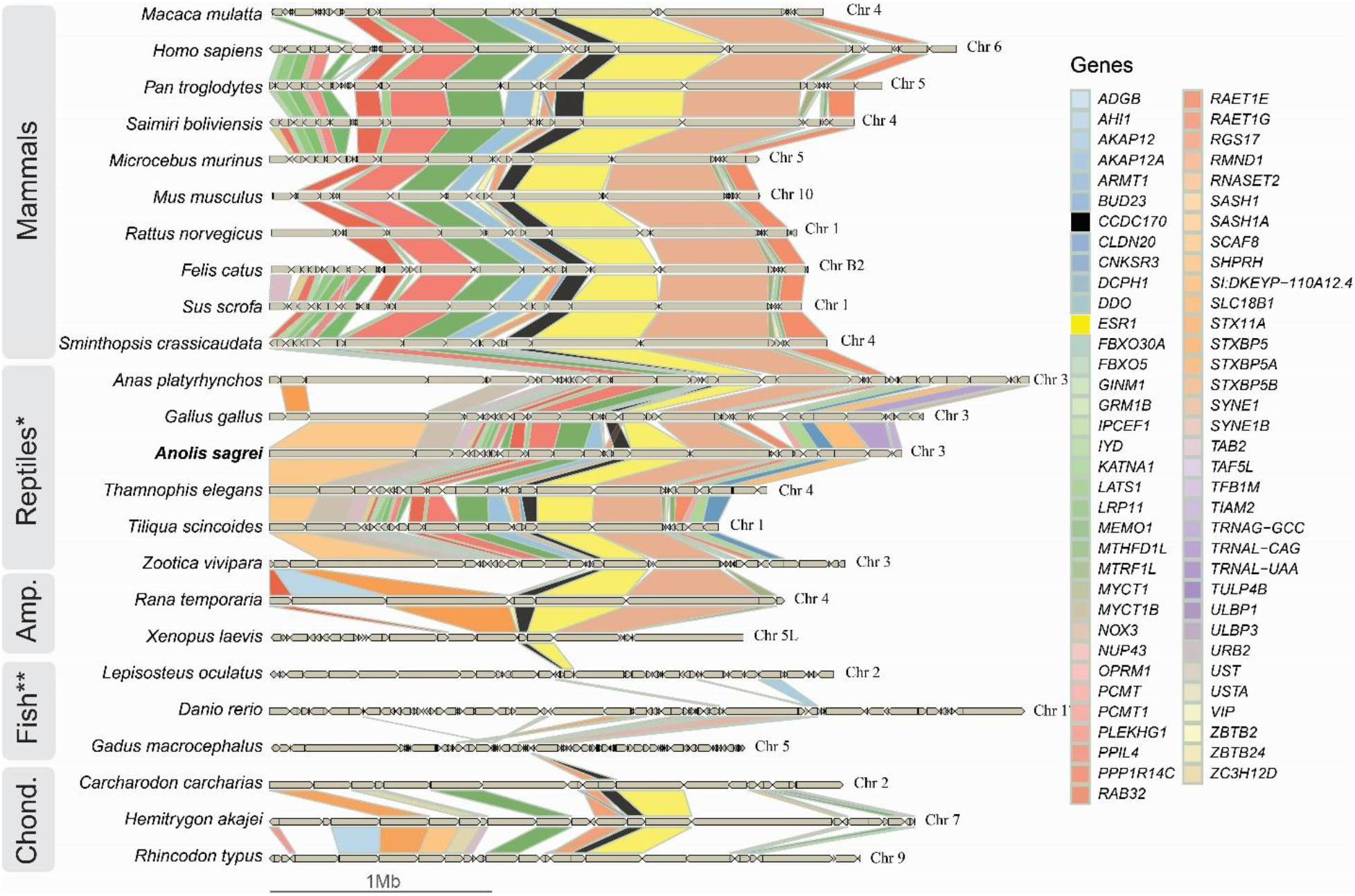
Conservation of the *CCDC170-ESR1*-locus in vertebrate genomes. Homology of genes in the *CCDC170-ESR1* surrounding region is shown by colored rectangles, and the direction of transcription is indicated by a triangle at the 3’-end. The focal species, *A. sagrei*, is shown in bold, and *CCDC170* is marked in black and *ESR1* in yellow. Representative species were selected from mammals, reptiles (* including birds), amphibians, fish (** actinopterygian fish) and chondrichthyans.

## Discussion

The female-limited color pattern polymorphism in *Anolis sagrei* has been mapped to the gene *CCDC170*, which is located directly upstream of the *ESR1* gene that is more highly expressed in female than in male embryos (Feiner et al., 2022). This genomic configuration provides a potential explanation for why the morph is sex-limited, and we propose a threshold model that captures why allelic variation at the color determining locus is masked in males as a consequence of low *ESR1* expression. In support of this model, we show that genetic male embryos treated with estradiol, causing them to express *ESR1* and *CCDC170* at levels similar to those of genetic females, developed color morphs in accordance with allelic variation at the *CCDC170* locus. Similarly, reducing expression of *ESR1* and *CCDC170* with fadrozole caused genetic females to develop a chevron morph even if they carry alleles for the diamond morph. These results suggest that expression levels of *CCDC170* are sensitive to estradiol levels, and that the naturally high levels of estradiol in female embryos causes the diamond-chevron polymorphism to be limited to females.

Our study does not uncover how exactly estradiol levels affect *CCDC170* expression and several mechanisms are plausible. First, *CCDC170* could be directly regulated by estradiol. Two predicted estrogen-response elements (EREs) are located in an intron of *CCDC170*, but it is unclear if they alone are able to upregulate the transcription of *CCDC170*. Second, *CCDC170* may be co-expressed with *ESR1*, which is regulated by a positive feedback loop that is responsive to estradiol enabled by the eight EREs located in the *ESR1* locus, three of which directly upstream. Physical proximity alone can partially explain co-expression (Ng et al., 2009; Zhao et al., 2026), but the two genes may also share a regulatory landscape with shared epigenetic features (e.g., DNA-methylation, histone modifications) and shared cis-regulatory elements. This idea is supported by the fact that co-expression between *ESR1* and *CCDC170* has been reported in humans (Dunbier et al., 2011) and was also found in our previous study (Feiner et al., 2022). A third plausible mechanism, though perhaps less likely, is that *CCDC170* is regulated via an unknown factor in *trans* that is itself regulated by estradiol. The fact that administration of estradiol and fadrozole leads to sex reversal (Ehl et al., 2017; Equinox, 2022) demonstrates the pervasive, systemic effect of these treatments and suggests that expression levels of a large number of genes are estradiol sensitive, as has been shown for mice, for example (Lacouture et al., 2025; Zhao et al., 2026). To ultimately discriminate between these three potential mechanisms of direct regulation of *CCDC170, cis*-regulation by *ESR1* or *trans*- regulation by an unknown factor, dissecting the transcriptional regulation of the *CCDC170-*(*ESR1*)*-*locus, or independently manipulating the *CCDC170* and *ESR1* genes are promising avenues for future research. Nevertheless, given the co-expression of *CCDC170* and *ESR1*, a *cis-*regulatory mechanism seems to be the most likely explanation for the estradiol-sensitivity of *CCDC170* and thus cause for the polymorphism to be female-limited.

Taking an evolutionary perspective, an intriguing question is whether this configuration with a co-location of a gene that regulates color pattern formation and a gene crucial for sex differentiation has been exploited by other lineages. To the best of our knowledge, in all vertebrates with a female-limited polymorphism for which the genetic basis has been resolved, the morph determining locus has been located on a sex chromosome (para guppy, Lindholm et al., 2004; common cuckoo, Merondun et al., 2024; Sandkam et al., 2021). Yet, the deep evolutionary conservation of the collinearity of *CCDC170* and *ESR1* across vertebrates (with the exception of actinopterygian fish) provides the possibility that the role of the *CCDC170-ESR1*-locus in regulating female-limited color pattern polymorphisms may be more widespread. Targeting the genetic basis of such polymorphisms in other *Anolis* species (Medina et al., 2016) or other vertebrates, such as the white-necked jacobin hummingbird (Falk et al., 2022) may reveal further instances. Taking a comparative approach to reconstruct the evolutionary steps that led up to the genetic architecture integrating morph determination and sex differentiation would provide insights into the mechanistic underpinnings of evolutionary transitions between sex-limited and universal, and between mono- and polymorphic color pattern systems.

While our study generally corroborates the causal role of *CCDC170* as the morph determining locus, we do not yet understand how allelic variation at this locus affects the migration, differentiation or other behaviors of pigment cells generate color patterns. In contrast to our previous work (Feiner et al., 2022), we did not detect consistent effects of treatment or sex on the expression of *CCDC170* in the tissue where pattern formation is taking place, the dorsal epithelia. Possible explanations may be that the experiment reported by Feiner et al. (2022) and those reported here used slightly different developmental stages or technical variation in the micro-dissection of the epithelial tissues, which resulted in overall lower expression levels in the present study. Pigment cells are embedded in various other cell types, which means that the extent to which other tissue layers are included could affect variation in qPCR results between studies. Application of single-cell RNA-sequencing or spatial transcriptomic techniques could overcome this problem by generating *CCDC170* and *ESR1* expression levels by cell type.

Our work demonstrates that morph determination is genetically controlled and sensitive to hormones, and, in addition, we also obtained important insights into the regulation of within-morph variation. Prompted by the observation that estradiol-treated males develop diamond morphs that are significantly more bar-like relative to those observed in control (ethanol-treated) females, we tested if also incubation temperature affects the quality of the expressed diamond pattern. Incubation temperatures outside natural ranges, in particular higher temperature (Angilletta, 2009; Pettersen et al., 2023), are known to cause developmental stress that can manifests in pervasive changes in global transcriptomic profiles (Feiner et al., 2018) as well as in specific gene expression changes associated with specific phenotypes, for example a shortened snout (Sanger et al., 2021). We found that stressfully high incubation temperatures gave rise to more bar-like diamond patterns relative to low or optimal temperatures. Together, these observations suggest that bar-like patterns may be a sign of developmental stress caused by high incubation temperatures or unnaturally high levels of estradiol. In line with this interpretation, we observed that ethanol-treated (control) females tended to show more bar-like patterns relative to untreated females incubated at optimal incubation temperatures (mid temperature in temperature incubation experiment), which leads us to speculate that not only high incubation temperature and unnaturally high levels of estradiol, but also ethanol may cause developmental stress. Unfortunately, our current understanding of the cellular developmental process of (diamond) pattern formation is too rudimentary to allow any speculation on how exactly this stress is reflected in cellular behaviors that manifest in bar-like rather than diamond-like patterns.

In conclusion, we present and experimentally validate a threshold model that captures the cross-talk between the color morph determination and sex differentiation systems to explain the female-limited diamond-chevron polymorphism in the brown anole lizard *Anolis sagrei*. We expose the co-location, and thereby co-regulation of the *CCDC170-ESR1*-locus as the mechanistic basis that allowed morph determination to be co-opted by the sex differentiation system, and thus provide insights into the developmental origins of color polymorphisms.

## Acknowledgement

We thank the animal caretakers at the Max Planck Institute in Plön, Germany, for assistance in the fadrozole experiments, Jane Jönsson at Lund University for guidance in the laboratory, Lukáš Kratochvíl for guidance on hormone treatments and Tobias Uller for feedback on the manuscript.

## Data and code availability statement

Data and code that was used to generate the results presented in this study are available at Github (accession number: *[will be added upon acceptance]*)

## Ethics approval statement

The study was conducted according to the Lund University Local Ethical Review Process, the Federation of European Laboratory Animal Science Associations guidelines and the German animal welfare law (Tierschutzgesetz § 11).

## References

Angilletta, M. J. (2009). Thermal Adaptation: A Theoretical and Empirical Synthesis.: Oxford University Press.

Bhatnagar, A. S., Häusler, A., Schieweck, K., Browne, L. J., Bowman, R., & Steele, R. E. (1990). Novel aromatase inhibitors. J Steroid Biochem Mol Biol, 37(3), 363–367. doi:10.1016/0960-0760(90)90485-4

Bondesson, M., Hao, R., Lin, C.-Y., Williams, C., & Gustafsson, J.-Å. (2015). Estrogen receptor signaling during vertebrate development. Biochimica et Biophysica Acta (BBA) - Gene Regulatory Mechanisms, 1849(2), 142–151. doi:10.1016/j.bbagrm.2014.06.005

Bull, J. J., Gutzke, W. H., & Crews, D. (1988). Sex reversal by estradiol in three reptilian orders. Gen Comp Endocrinol, 70(3), 425–428. doi:10.1016/0016-6480(88)90117-7

Cardone, A., Angelini, F., & Varriale, B. (1998). Autoregulation of Estrogen and Androgen Receptor mRNAs and Downregulation of Androgen Receptor mRNA by Estrogen in Primary Cultures of Lizard Testis Cells. General and Comparative Endocrinology, 110(3), 227–236. doi:10.1006/gcen.1998.7063

Chen, G., Zhou, T., Chen, M., Zou, G., & Liang, H. (2022). Effect of Estradiol on Estrogen Nuclear Receptors Genes Expression on Embryonic Development Stages in Chinese Soft-Shelled Turtle (Pelodiscus sinensis). Fishes, 7(5), 223. Retrieved from doi:10.3390/fishes7050223

Dijkstra, P. D., Seehausen, O., & Groothuis, T. G. G. (2007). Intrasexual competition among females and the stabilization of a conspicuous colour polymorphism in a Lake Victoria cichlid fish. Proceedings of the Royal Society B: Biological Sciences, 275(1634), 519–526. doi:10.1098/rspb.2007.1441

Dunbier, A. K., Anderson, H., Ghazoui, Z., Lopez-Knowles, E., Pancholi, S., Ribas, R., … Dowsett, M. (2011). ESR1 is co-expressed with closely adjacent uncharacterised genes spanning a breast cancer susceptibility locus at 6q25.1. PLoS Genet, 7(4), e1001382. doi:10.1371/journal.pgen.1001382

Ehl, J., Vukić, J., & Kratochvíl, L. (2017). Hormonal and thermal induction of sex reversal in the bearded dragon (Pogona vitticeps, Agamidae). Zoologischer Anzeiger, 271, 1–5. doi:10.1016/j.jcz.2017.11.002

Equinox, B. M. (2022). Steroid hormones, architects of behavior and development: a look at the effects of sex steroid hormones in a lizards model.. (Doctor of Philosophy), University of Oklahoma, Oklahoma.

Falk, J. J., Rubenstein, D. R., Rico-Guevara, A., & Webster, M. S. (2022). Intersexual social dominance mimicry drives female hummingbird polymorphism. Proc Biol Sci, 289(1982), 20220332. doi:10.1098/rspb.2022.0332

Feiner, N., Brun-Usan, M., Andrade, P., Pranter, R., Park, S., Menke, D. B., … Uller, T. (2022). A single locus regulates a female-limited color pattern polymorphism in a reptile. Science Advances, 8(10), eabm2387. doi:10.1126/sciadv.abm2387

Feiner, N., Munch, K. L., Jackson, I. S. C., & Uller, T. (2020). Enhanced locomotor performance on familiar surfaces is uncoupled from morphological plasticity in Anolis lizards. J Exp Zool A Ecol Integr Physiol. doi:10.1002/jez.2349

Feiner, N., Rago, A., While, G. M., & Uller, T. (2018). Developmental plasticity in reptiles: Insights from temperature-dependent gene expression in wall lizard embryos. J Exp Zool A Ecol Integr Physiol, 329(6-7), 351–361. doi:10.1002/jez.2175

Gamble, T., & Zarkower, D. (2014). Identification of sex-specific molecular markers using restriction site-associated DNA sequencing. Mol Ecol Resour, 14(5), 902–913. doi:10.1111/1755-0998.12237

Geneva, A. J., Park, S., Bock, D. G., de Mello, P. L. H., Sarigol, F., Tollis, M., … Losos, J. B. (2022). Chromosome-scale genome assembly of the brown anole (Anolis sagrei), an emerging model species. Communications Biology, 5(1), 1126. doi:10.1038/s42003-022-04074-5

Hendrickx, F., De Corte, Z., Sonet, G., Van Belleghem, S. M., Köstlbacher, S., & Vangestel, C. (2022). A masculinizing supergene underlies an exaggerated male reproductive morph in a spider. Nature Ecology & Evolution, 6(2), 195–206. doi:10.1038/s41559-021-01626-6

Jiang, P., Li, Y., Poleshko, A., Medvedeva, V., Baulina, N., Zhang, Y., … Chen, X. (2017). The Protein Encoded by the CCDC170 Breast Cancer Gene Functions to Organize the Golgi-Microtubule Network. EBioMedicine, 22, 28–43. doi:10.1016/j.ebiom.2017.06.024

Katsu, Y., Matsubara, K., Kohno, S., Matsuda, Y., Toriba, M., Oka, K., … Iguchi, T. (2010). Molecular cloning, characterization, and chromosome mapping of reptilian estrogen receptors. Endocrinology, 151(12), 5710–5720. doi:10.1210/en.2010-0356

Kratochwil, C. F., & Mallarino, R. (2023). Mechanisms Underlying the Formation and Evolution of Vertebrate Color Patterns. Annu Rev Genet, 57, 135–156. doi:10.1146/annurev-genet-031423-120918

Kuriyama, T., Murakami, A., Brandley, M., & Hasegawa, M. (2020). Blue, Black, and Stripes: Evolution and Development of Color Production and Pattern Formation in Lizards and Snakes. Frontiers in Ecology and Evolution, Volume 8-2020. doi:10.3389/fevo.2020.00232

Lacouture, A., Sylla, M. S., Germain, L., & Audet-Walsh, É. (2025). RNA-seq dataset of the estrogen-dependent regulation of the transcriptome in mouse mammary gland organoids. Data in Brief, 62, 111984. doi:10.1016/j.dib.2025.111984

Lindholm, A. K., Brooks, R., & Breden, F. (2004). Extreme polymorphism in a Y-linked sexually selected trait. Heredity, 92(3), 156–162. doi:10.1038/sj.hdy.6800386

Lung, D. K., Reese, R. M., & Alarid, E. T. (2020). Intrinsic and Extrinsic Factors Governing the Transcriptional Regulation of ESR1. Hormones and Cancer, 11(3), 129–147. doi:10.1007/s12672-020-00388-0

Mank, J. E. (2023). Sex-specific morphs: the genetics and evolution of intra-sexual variation. Nature Reviews Genetics, 24(1), 44–52. doi:10.1038/s41576-022-00524-2

Marlatt, V. L., Martyniuk, C. J., Zhang, D., Xiong, H., Watt, J., Xia, X., … Trudeau, V. L. (2008). Auto-regulation of estrogen receptor subtypes and gene expression profiling of 17beta-estradiol action in the neuroendocrine axis of male goldfish. Mol Cell Endocrinol, 283(1-2), 38–48. doi:10.1016/j.mce.2007.10.013

Medina, I., Losos, J. B., & Mahler, D. L. (2016). Evolution of dorsal pattern variation in Greater Antillean Anolis lizards. Biological Journal of the Linnean Society, n/a(n/a). doi:10.1111/bij.12881

Merondun, J., Marques, C. I., Andrade, P., Meshcheryagina, S., Galván, I., Afonso, S., … Wolf, J. B. W. (2024). Evolution and genetic architecture of sex-limited polymorphism in cuckoos. Science Advances, 10(17), eadl5255. doi:10.1126/sciadv.adl5255

Moon, R. M., & Kamath, A. (2019). Re-examining escape behaviour and habitat use as correlates of dorsal pattern variation in female brown anole lizards, Anolis sagrei (Squamata: Dactyloidae). Biological Journal of the Linnean Society, 126(4), 783–795. doi:10.1093/biolinnean/blz006

Ng, Y. K., Wu, W., & Zhang, L. (2009). Positive correlation between gene coexpression and positional clustering in the zebrafish genome. BMC Genomics, 10, 42. doi:10.1186/1471-2164-10-42

Nishikawa, H., Iijima, T., Kajitani, R., Yamaguchi, J., Ando, T., Suzuki, Y., … Fujiwara, H. (2015). A genetic mechanism for female-limited Batesian mimicry in Papilio butterfly. Nature Genetics, 47(4), 405–409. doi:10.1038/ng.3241

Pearson, P., & Warner, D. (2016). Habitat- and season-specific temperatures affect phenotypic development of hatchling lizards. Biology Letters, 12, 20160646. doi:10.1098/rsbl.2016.0646

Pettersen, A. K., Feiner, N., Noble, D. W. A., While, G. M., Uller, T., & Cornwallis, C. K. (2023). Maternal behavioral thermoregulation facilitated evolutionary transitions from egg laying to live birth. Evol Lett, 7(5), 351–360. doi:10.1093/evlett/qrad031

Rice, E. S., Kohno, S., John, J. S., Pham, S., Howard, J., Lareau, L. F., … Green, R. E. (2017). Improved genome assembly of American alligator genome reveals conserved architecture of estrogen signaling. Genome Res, 27(5), 686–696. doi:10.1101/gr.213595.116

Saceda, M., Lippman, M. E., Chambon, P., Lindsey, R. L., Ponglikitmongkol, M., Puente, M., & Martin, M. B. (1988). Regulation of the Estrogen Receptor in MCF-7 Cells by Estradiol. Molecular Endocrinology, 2(12), 1157–1162. doi:10.1210/mend-2-12-1157

Sandkam, B. A., Almeida, P., Darolti, I., Furman, B. L. S., van der Bijl, W., Morris, J., … Mank, J. E. (2021). Extreme Y chromosome polymorphism corresponds to five male reproductive morphs of a freshwater fish. Nature Ecology & Evolution, 5(7), 939–948. doi:10.1038/s41559-021-01452-w

Sanger, T. J., Harding, L., Kyrkos, J., Turnquist, A. J., Epperlein, L., Nunez, S. A., … Czesny, B. (2021). Environmental Thermal Stress Induces Neuronal Cell Death and Developmental Malformations in Reptiles. Integr Org Biol, 3(1), obab033. doi:10.1093/iob/obab033

Sanger, T. J., Kyrkos, J., Lachance, D. J., Czesny, B., & Stroud, J. T. (2018). The effects of thermal stress on the early development of the lizard Anolis sagrei. J Exp Zool A Ecol Integr Physiol, 329(4-5), 244–251. doi:10.1002/jez.2185

Sanger, T. J., Losos, J. B., & Gibson-Brown, J. J. (2008). A developmental staging series for the lizard genus Anolis: a new system for the integration of evolution, development, and ecology. J Morphol, 269(2), 129–137. doi:10.1002/jmor.10563

Shine, R., Warner, D. A., & Radder, R. (2007). Windows of embryonic sexual lability in two lizard species with environmental sex determination. Ecology, 88(7), 1781–1788. doi:10.1890/06-2024.1

Shupnik, M. A., Gordon, M. S., & Chin, W. W. (1989). Tissue-specific regulation of rat estrogen receptor mRNAs. Mol Endocrinol, 3(4), 660–665. doi:10.1210/mend-3-4-660

Zhao, Y., Liu, R., Ng, J. P., Yu, S., Ahn, I. S., Diamante, G., … Yang, X. (2026). Estradiol treatment induces both shared and unique gene regulation and networks in adipose cell types of gonadectomized obese XX and XY mice. Biol Sex Differ, 17(1). doi:10.1186/s13293-026-00859-z

