## Supplementary Table 1 and 2 for "Towards understanding the mechanistic basis of a sex-limited color polymorphism"

This document contains 2 Supplementary Tables.

Supplementary material

**Supplementary Table S1.** List of reagents used in this study.

| Reagent | Company | Catalog# | Cas# |
|---|---|---|---|
| DNeasy Blood & Tissue Kit | Qiagen | 69504 | |
| GoTaq Hot Start Master Mix | Promega | M5122 | |
| TaqI-v2 | New England Biolabs | R0149S | |
| rCutSmart Buffer | New England Biolabs | B6004S | |
| RNeasy Micro Kit | Qiagen | 74004 | |
| 5x First strand buffer | Invitrogen | Y02321 | |
| Dithiothreitol (DTT) | Invitrogen | Y00147 | |
| SupersScript III reverse transcriptase | Invitrogen | 18080 | |
| RNasin Plus RNase Inhibitor | Promega | N261A | |
| RNase H | New England Biolabs | M0297S | |
| Platinum SYBR Green qPCR SuperMix-UDG | Thermo Fisher Scientific | 11733046 | |
| ROX | Invitrogen | 54881 | 50-28-2 |
| Phosphate buffered saline (PBS) | Sigma-Aldrich | P4417 | 7732-18-5 |
| Water, nucl. free, Mol. Biol. Grade, Ultrapure | ThermoScientific | J71786.K8 | |
| 17β-Estradiol | Cayman chemical | 10006315 | 102676-31-3 |
| Fadrozole hydrochloride | Sigma-Aldrich | F3806 | |
| RNAlater Solution | Invitrogen | AM70 | |

Supplementary material

**Supplementary Table S2.** Genomes and genome annotations used for collinearity analysis.

| File | Species | FTP | Accession |
|---|---|---|---|
| GCF_000001635.27_GRCm39_genomic.gff | *Mus musculus* | https://ftp.ncbi.nlm.nih.gov/genomes/all/GCF/000/001/635/GCF_000001635.27_GRCm39/ | GCF_000001635.27 |
| GCF_000003025.6_Sscrofa11.1_genomic.gff | *Sus scrofa* | https://ftp.ncbi.nlm.nih.gov/genomes/all/GCF/000/003/025/GCF_000003025.6_Sscrofa11.1/ | GCF_000003025.6 |
| GCF_018350175.1_F.catus_Fca126_mat1.0_genomic.gff | *Felis catus* | https://ftp.ncbi.nlm.nih.gov/genomes/all/GCF/018/350/175/GCF_018350175.1_F.catus_Fca126_mat1.0/ | GCF_018350175.1 |
| GCF_035046505.1_rTilSci1.hap2_genomic.gff | *Tiliqua scincoides* | https://ftp.ncbi.nlm.nih.gov/genomes/all/GCF/035/046/505/GCF_035046505.1_rTilSci1.hap2/ | GCF_035046505.1 |
| GCF_009914755.1_T2T-CHM13v2.0_genomic.gff | *Homo sapiens* | https://ftp.ncbi.nlm.nih.gov/genomes/all/GCF/009/914/755/GCF_009914755.1_T2T-CHM13v2.0/ | GCF_009914755.1 |
| GCF_016699485.2_bGalGal1.mat.broiler.GRCg7b_genomic.gff | *Gallus gallus* | https://ftp.ncbi.nlm.nih.gov/genomes/all/GCF/016/699/485/GCF_016699485.2_bGalGal1.mat.broiler.GRCg7b/ | GCF_016699485.2 |
| GCF_017639515.1_sCarCar2.pri_genomic.gff | *Carcharodon carcharias* | https://ftp.ncbi.nlm.nih.gov/genomes/all/GCF/017/639/515/GCF_017639515.1_sCarCar2.pri/ | GCF_017639515.1 |

| | | | |
|---|---|---|---|
| GCF_017654675.1_Xenopus_laevis_v10.1_genomic.gff | *Xenopus laevis* | https://ftp.ncbi.nlm.nih.gov/genomes/all/GCF/017/654/675/GCF_017654675.1_Xenopus_laevis_v10.1/ | GCF_017654675.1 |
| GCF_028858775.2_NHGRI_mPanTro3-v2.0_pri_genomic.gff | *Pan troglodytes* | https://ftp.ncbi.nlm.nih.gov/genomes/all/GCF/028/858/775/GCF_028858775.2_NHGRI_mPanTro3-v2.0_pri/ | GCF_028858775.2 |
| GCF_031168955.1_ASM3116895v1_genomic.gff | *Gadus macrocephalus* | https://ftp.ncbi.nlm.nih.gov/genomes/all/GCF/031/168/955/GCF_031168955.1_ASM3116895v1/ | GCF_031168955.1 |
| GCF_036323735.1_GRCr8_genomic.gff | *Rattus norvegicus* | https://ftp.ncbi.nlm.nih.gov/genomes/all/GCF/036/323/735/GCF_036323735.1_GRCr8/ | GCF_036323735.1 |
| GCF_037176765.1_rAnoSag1.mat_genomic.gff | *Anolis sagrei* | https://ftp.ncbi.nlm.nih.gov/genomes/all/GCF/037/176/765/GCF_037176765.1_rAnoSag1.mat/ | GCF_037176765.1 |
| GCF_040939455.1_M.murinus_Inina_mat1.0_genomic.gff | *Microcebus murinus* | https://ftp.ncbi.nlm.nih.gov/genomes/all/GCF/040/939/455/GCF_040939455.1_M.murinus_Inina_mat1.0/ | GCF_040939455.1 |
| GCF_047663525.1_IASCAAS_PekinDuck_T2T_genomic.gff | *Anas platyrhynchos* | https://ftp.ncbi.nlm.nih.gov/genomes/all/GCF/047/663/525/GCF_047663525.1_IASCAAS_PekinDuck_T2T/ | GCF_047663525.1 |
| GCF_048418815.1_sHemAka1.3_genomic.gff | *Hemitrygon akajei* | https://ftp.ncbi.nlm.nih.gov/genomes/all/GCF/048/418/815/GCF_048418815.1_sHemAka1.3/ | GCF_048418815.1 |

| | | | |
|---|---|---|---|
| GCF_021869965.1_sRhiTyp1.1_genomic.gff | *Rhincodon typus* | https://ftp.ncbi.nlm.nih.gov/genomes/all/GCF/021/869/965/GCF_021869965.1_sRhiTyp1.1/ | GCF_021869965.1 |
| GCF_048565385.1_mSaiBol1.pri_genomic.gff | *Saimiri boliviensis* | https://ftp.ncbi.nlm.nih.gov/genomes/all/GCF/048/565/385/GCF_048565385.1_mSaiBol1.pri/ | GCF_048565385.1 |
| GCF_048593235.1_ASM4859323v1_genomic.gff | *Sminthopsis crassicaudata* | https://ftp.ncbi.nlm.nih.gov/genomes/all/GCF/048/593/235/GCF_048593235.1_ASM4859323v1/ | GCF_048593235.1 |
| GCF_049306965.1_GRCz12tu_genomic.gff | *Danio rerio* | https://ftp.ncbi.nlm.nih.gov/genomes/all/GCF/049/306/965/GCF_049306965.1_GRCz12tu/ | GCF_049306965.1 |
| GCF_040954835.1_fLepOcu1.hap2_genomic.gff | *Lepisosteus oculatus* | https://ftp.ncbi.nlm.nih.gov/genomes/all/GCF/040/954/835/GCF_040954835.1_fLepOcu1.hap2/ | GCF_040954835.1 |
| GCF_049350105.2_T2T-MMU8v2.0_genomic.gff | *Macaca mulatta* | https://ftp.ncbi.nlm.nih.gov/genomes/all/GCF/049/350/105/GCF_049350105.2_T2T-MMU8v2.0/ | GCF_049350105.2 |
| GCF_905171775.1_aRanTem1.1_genomic.gff | *Rana temporaria* | https://ftp.ncbi.nlm.nih.gov/genomes/all/GCF/905/171/775/GCF_905171775.1_aRanTem1.1/ | GCF_905171775.1 |
| GCF_963506605.1_rZooViv1.1_genomic.gff | *Zootica vivipara* | https://ftp.ncbi.nlm.nih.gov/genomes/all/GCF/963/506/605/GCF_963506605.1_rZooViv1.1/ | GCF_963506605.1 |

| | | | |
|---|---|---|---|
| GCF_009769535.1_rThaEle1.pri_genomic.gff | *Thamnophis elegans* | https://ftp.ncbi.nlm.nih.gov/genomes/all/GCF/009/769/535/GCF_009769535.1_rThaEle1.pri/ | GCF_009769535.1 |
